# dartR v2: an accessible genetic analysis platform for conservation, ecology, and agriculture

**DOI:** 10.1101/2022.03.30.486475

**Authors:** Jose Luis Mijangos, Bernd Gruber, Oliver Berry, Carlo Pacioni, Arthur Georges

## Abstract

1. Innumerable approaches to analyse genetic data are now available to guide conservation, ecological and agricultural projects. However, streamlined and accessible tools are needed to bring these approaches within reach of a broader user base. dartR was released in 2018 to lessen the intrinsic complexity of analysing single nucleotide polymorphisms (SNPs) and dominant markers (presence/absence of amplified sequence tags) by providing user-friendly data quality control and marker selection functions. dartR users have grown steadily since its release and provided valuable feedback on their interaction with the package allowing us to enhance dartR capabilities.
2. Here, we present Version 2 of dartR. In this iteration, we substantially increased the number of available functions from 45 to 144. In addition to improved functionality, we focused on enhancing the user experience by extending plot customisation, function standardisation, increasing user support and function speed. dartR provides functions for various stages in analysing genetic data, from data manipulation to reporting.
3. dartR provides many functions for importing, exporting and linking to other packages, to provide an easy-to-navigate conduit between data generation and analysis options already available via other packages. We also implemented simulation functions whose results can be analysed seamlessly with several other dartR functions.
4. As more methods and approaches mature to inform conservation, we envision that accessible platforms to analyse genetic data will play a crucial role in translating science into practice.

## Introduction

The plummeting costs of DNA sequencing have opened a powerful window of opportunity to use genetic data to inform biodiversity conservation, restoration of ecosystems, invasive species management and breeding of animals and plants (Breed *et al*., 2019). Remarkably, applied genetic studies have transitioned from typically analysing a dozen molecular markers to tens and even hundreds of thousands of markers in less than a decade. Similarly, the process of marker development that could take months of laboratory work a decade ago has been taken over by sequencing companies using novel approaches, such as genotyping by sequencing (Narum et al., 2013) or using restriction enzymes to reduce genome complexity (DArTseq; Kilian et al., 2012). These technological advances are reflected in the growing number and diversity of ways genetic data is analysed and applied. (*e.g*. identification of adaptive variation is now within reach for non-model organisms; Weigand & Leese, 2018)

Even though genetic data are increasingly accessible and population genomics has proved to be a powerful tool to improve biodiversity conservation and ecological restoration efforts (Garner *et al*., 2016; Hohenlohe *et al*., 2021), genetic information is not yet regularly used outside of the research community (Shafer et al., 2015). BSeveral barriers to bridging this gap between research and practice have been identified, including poor communication between researchers and other stakeholders, insufficient funding, and lack of genetics expertise (Taylor et al., 2017). A further barrier is arguably the intrinsic complexity involved in analysing genetic data. For instance, to interpret analysis results appropriately, it is necessary to understand theoretical models and population genetics principles (Andrews & Luikart, 2014). Furthermore, advanced computer and programming skills and the use of several programs, which are often complex and time-consuming to master, are required to make full use of the genetic data (Hohenlohe et al., 2021). Therefore, today, it is no longer the time needed for DNA sequencing that limits the speed of results, but rather a deficit of knowledge and skills to analyse genetic data.

dartR, an R package for analysing single nucleotide polymorphisms (SNPs) and presence/absence of amplified sequence tags was released in 2018 (Gruber et al., 2018) and designed to bridge the gap between science and practice. dartR aims to bring the timeframe to analyse genetic data into line with the timeframe required by stakeholders to make their decisions and at the same time provide a broad range of analyses and pipelines in a user-friendly platform that allows no programming expertise to do so. dartR leverages the capabilities of the open-source programming language R (R Core Team, 2021) and the robustness of the genlight object from the package adegenet for representing large genetic datasets (Jombart & Ahmed, 2011). In the four years since its release, dartR has grown a large user base, evidenced by several hundred daily downloads and an active Google group (https://groups.google.com/g/dartr). With the genomic revolution well underway, there is a constant and rapid diversification of new methods and analyses, which users seek to include in their work, ideally without switching between platforms.

Here we present a significant update of dartR. Our purpose is to bring diverse and sophisticated analytical tools within reach of a broad user base of genomic data. dartR facilitates all stages in analysing genetic data, from data quality control to the preparation of publishing quality plots through streamlined and accessible functions and strong user support, including tutorials, detailed function documentation, and error checking.

## What is new in dartR 2.0?

In dartR 2.0, we have added 99 functions to the initial 45 functions from version 1 (Fig. 1 and Supplementary Table 1). In response to user feedback, we provide users with a deeper understanding of the purpose of each function, its underlying theory and its limitations by expanding and improving our tutorials and function documentation. Additionally, we have implemented messages to communicate errors, warnings, reports, and important information while running each function. All the functions have been extensively tested, debugged, standardised, and their speed has been increased in many cases. Following the adage “a picture is worth a thousand words”, we have improved all the graphical outputs by standardising their format, increasing readability, and extending their scope for customisation.

**Figure 1.**
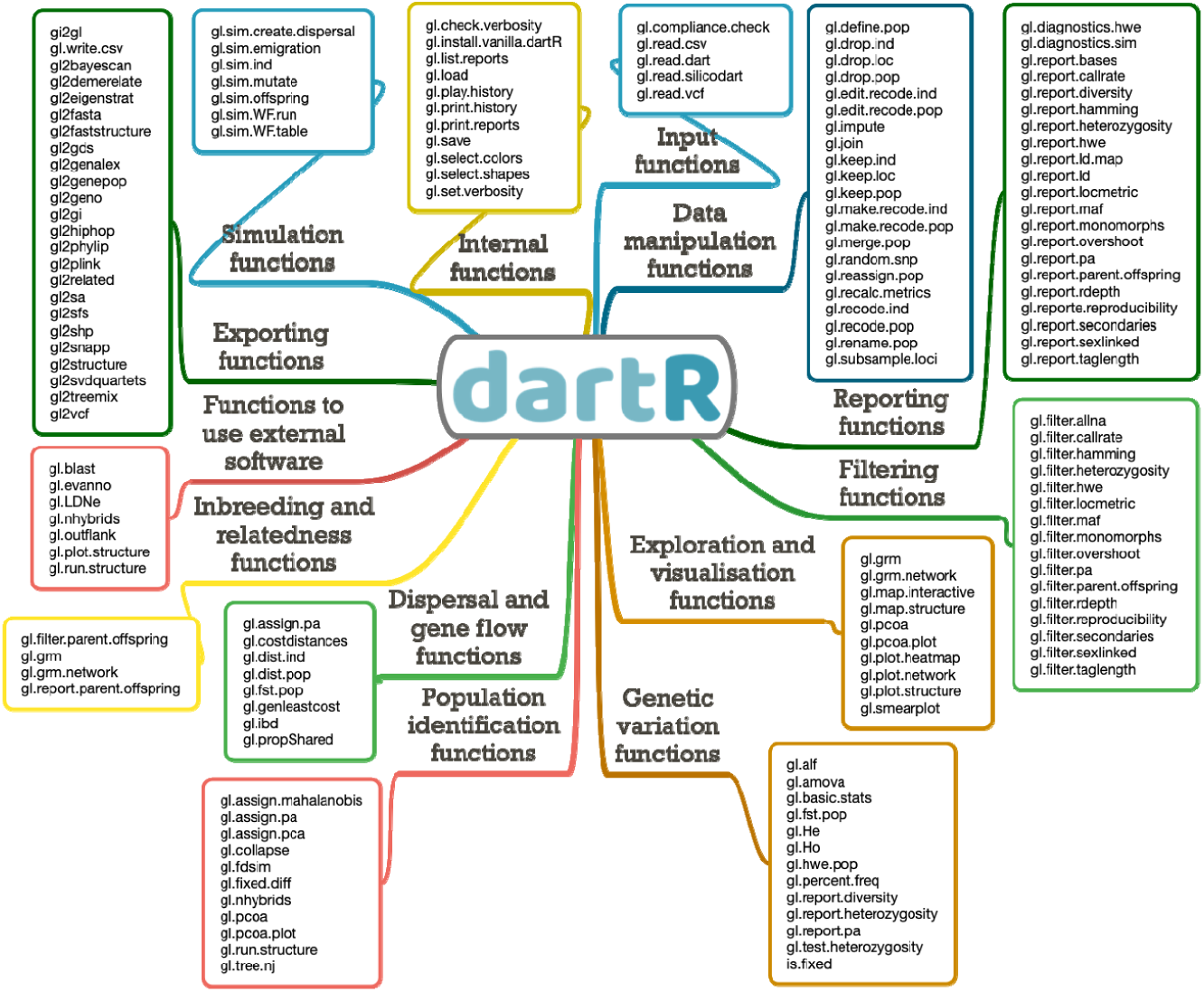
Overview of the functions currently available in dartR covering various stages in the analysis of genetic data.

We realised that many individual researchers had developed their own scripts and analyses, which would be very helpful for others if made available. Therefore, we encourage these “independent developers” to collaborate with dartR having provided a framework on how to write and document functions for dartR. To further encourage this collaboration, we have regularly developer meetings and personal support to integrate analyses of independent developers.

Initially, dartR aimed to primarily analyse the genomic data format provided by the sequencing company Diversity Arrays Technology Pty Ltd (DArT https://www.diversityarrays.com/). In version two, we extended dartR’s capabilities to import from and export to several formats to store SNP data to make dartR accessible to a broader pool of users.

## Function categories available in dartR

To facilitate the usage and identification of the resources available in dartR, we categorised the functions based on the different stages in the analysis of genetic data. Typical steps are data input, data manipulation, filtering, reporting, exploration, visualization, and analysis. We also provide tutorials to guide the user for the most relevant stages, which can be accessed at http://georges.biomatix.org/dartR. In this section, we enumerate dartR function categories while highlighting representative functions from each category.

As our basic format to **input** and store genetic data, we adopted the genlight object from the package adegenet (Jombart & Ahmed, 2011). One of the main attributes of the genlight object is its efficient data compression using a bit-level coding scheme. We extended the genlight object by adding two additional compartments containing metadata for individuals (ind.metrics) and loci (loc.metrics). dartR can read common formats, including FASTA, VCF, PLINK, DArTseq™, genepop and CSV files. To ensure the compatibility of the imported data, we developed the function **gl.compliance.check()** to inspect the elements within the genlight object and, if necessary, correct incompatibilities.

dartR offers functions to facilitate **data manipulation** for loci, individuals and populations, such as renaming individuals, assigning and reassigning them to populations, removing individuals, populations and loci, merging populations and subsampling individuals and loci. After data manipulation, some locus metrics will no longer apply; the function **gl.recalc.metrics()** will recalculate the various locus metrics as necessary.

The **filtering** process is a decisive step in analysing genetic data that depends on sensible threshold decisions (O’Leary *et al*., 2018). With this in mind, we provide a complementary reporting function for each of our 16 filtering functions. **Reporting** functions present the data in the form of summary statistics, tabulation of quantiles, boxplots, and histograms. In a two-stage process, users can use the results of reporting functions to implement thresholds in filter functions that are appropriate for their application and data characteristics. For example, identifying and filtering loci that deviate from Hardy-Weinberg proportions is essential in many workflows. Several technical and biological phenomena can cause this deviation and must be distinguished for correct interpretation of the data (Waples, 2015). Our functions **gl.diagnostics.hwe(), gl.report.hwe()** and **gl.filter.hwe()** allow the diagnosis, evaluation and filtering of loci deviating from Hardy-Weinberg proportions using either the Exact or the Chi-square method, adjustment for multiple comparisons and ternary plots (Fig. 2).

**Figure 2.**
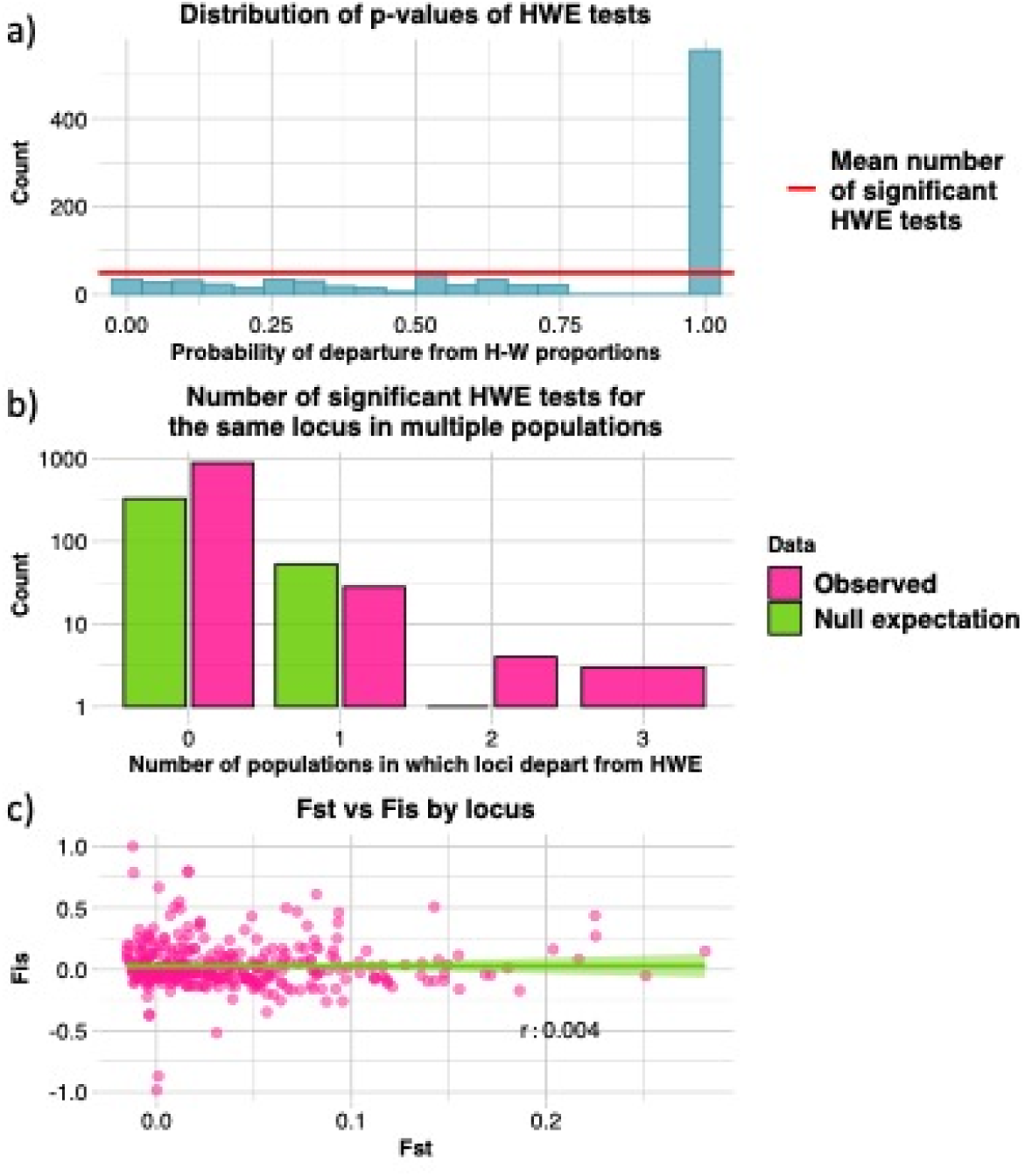
Output from function gl.diagnostics.hwe() which implements the recommendations from Waples (2015) and De Meeûs *et al* (2007). **a)** Histogram showing the distribution of p-values of Hardy-Weinberg Equilibrium (HWE) tests. The distribution should be roughly uniform across equal-sized bins. **b)** Bar plot showing observed and expected number of significant HWE tests for the same locus in multiple populations. If HWE tests are significant by chance alone, observed and expected number of HWE tests should have roughly a similar distribution. **c)** Scatter plot with a linear regression between *F*_*ST*_ and *F*_*IS*_, averaged across subpopulations. In the lower right corner of the plot, the Pearson correlation coefficient is reported. A positive relationship is expected in case of the presence of null alleles (De Meeûs, 2018).

The **exploration and visualisation** stage is critical to identify and interpret genetic patterns, generate hypotheses and set the path for downstream analyses. Functions for this stage in dartR include **gl.pcoa()** and **gl.pcoa.plot()**, which perform and plot principal component analysis (PCA; Fig. 3) and the related principal coordinates analysis (PCoA). PCA and PCoA are particularly suitable for genetic data. Despite not relying on genetic principles or models, results can reveal spatial patterns, evolutionary or ecological processes such as migration, geographical and reproductive isolation, and admixture (McVean, 2009). Other visualisation and exploration tools available include heatmaps, network plots, smear plots and mapping of sampling locations.

**Figure 3.**
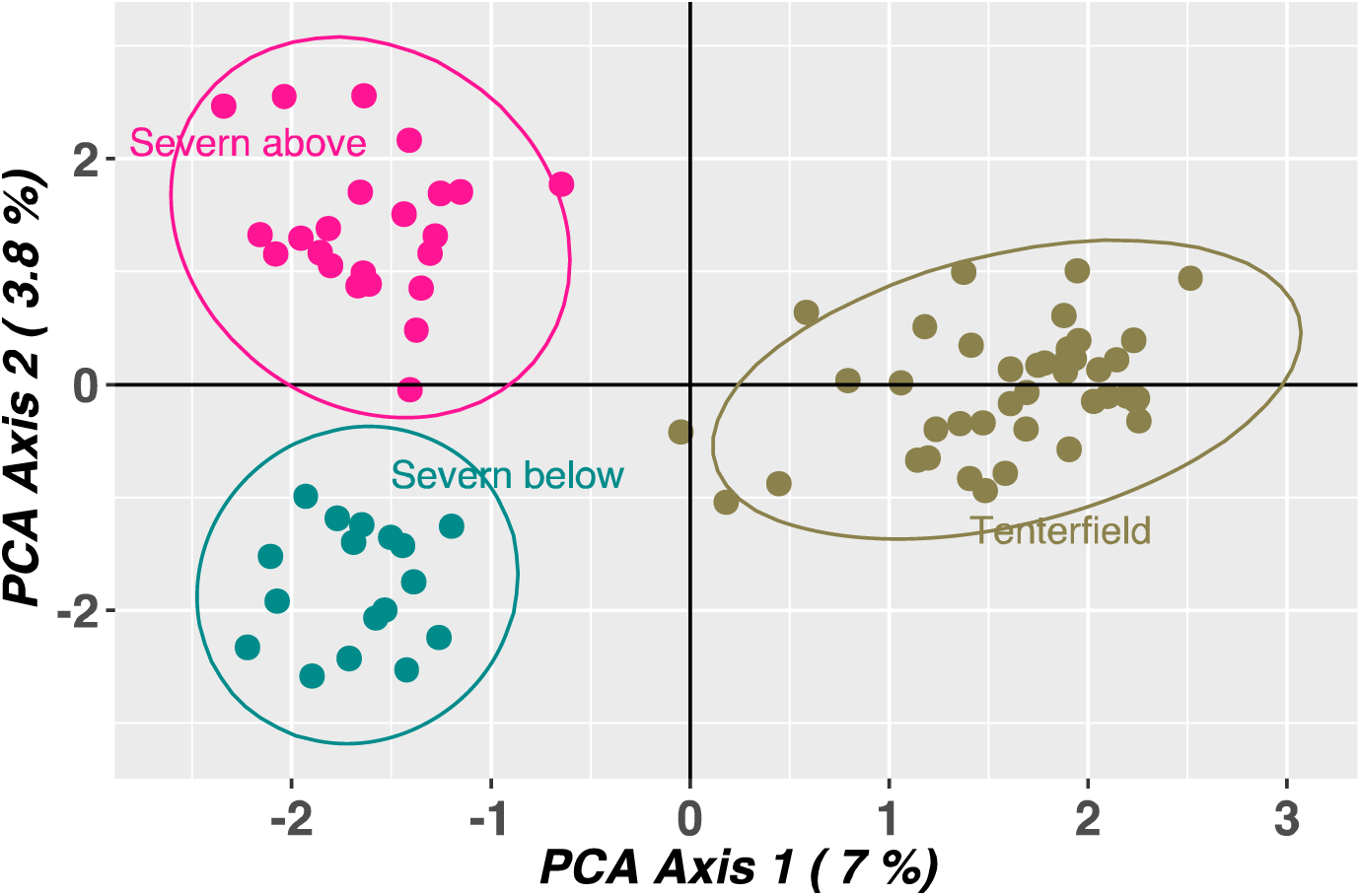
Principal component analyses (PCA) using a platypus dataset provided with the package. PCA shows that platypuses sampled below (Severn below) and above (Severn above) a large dam form separated clusters in contrast to platypuses sampled in an unregulated river (Tenterfield Creek).

Once the dartR user has read, manipulated, filtered and explored their genetic data, many **analyses** can be performed to inform the decision making, evaluation and monitoring processes of conservation, restoration and breeding projects. Genetic data can provide insights into biological processes on two different but tightly linked fronts: a) issues associated with genetic diversity and its relationship with fitness, such as inbreeding depression and evolutionary potential, and; b) demographic issues, such as dispersal, population size and hybridisation. dartR offers various functions that address both of these suites of processes.

*Genetic variation* can be monitored or evaluated with the function **gl.report.diversity()**, which calculates the q-profile, a spectrum of measures whose contrasting properties provide a rich summary of diversity, including allelic richness, Shannon information and heterozygosity (Sherwin et al., 2017). These measures are then converted to a standard scale of effective numbers (Hill’s numbers), so they can be directly compared. Other functions allow different aspects and metrics of diversity to be characterised by partitioning variation geographically using Analysis of Molecular Variance (AMOVA), statistical testing of heterozygosity difference between populations, or standardising heterozygosity estimates using the number of invariant sites.

*Identifying natural aggregations of individuals and populations* using genetic data has been an important tool to maximise and prioritise available resources in conservation and restoration projects, for example, to define evolutionarily significant units (ESUs; Funk et al., 2012), to delimitate species (Georges et al., 2018; Unmack et al., 2022), to identify populations suitable for eradication (Robertson & Gemmell, 2004) and to demarcate seed transfer zones for ecological restoration (Durka et al., 2017). dartR functions suitable for these applications include **gl.fixed.diff()**, which generates a matrix of fixed allelic differences between populations. The function **gl.collapse()** can be used to iteratively combine populations and aggregations of populations based on the absence of fixed allelic differences to yield a set of diagnosable units. These functions accommodate the risk of false positive fixed differences likely to occur when samples sizes are small. A further application of identifying populations is the assignment of individuals of unknown provenance to their source population, which is particularly important in wildlife forensics to support law enforcement (Bourret et al., 2020). Functions such as **gl.assign.pa()** and **gl.assign.pca()** are capable of assigning individuals of unknown provenance to a population using private alleles (*i.e*., alleles that are exclusive to particular populations) and standardized proximity, respectively.

*Dispersal and gene flow* are fundamental evolutionary and ecological processes that enable individuals to recolonise new habitat and replenish population’s gene pool (Tigano & Friesen, 2016). These processes can be investigated by assessing the correlation between genetic distance among populations or individuals and the geographic distance separating them (Cayuela et al., 2018). The function **gl.genleastcost()** performs a least-cost path analysis based on a friction matrix to test the hypothesis that genetic distance correlates with landscape attributes, such as barriers or habitat corridors, rather than geographic distance. Other functions include the calculation of several genetic distances between individuals and populations, testing for isolation by distance (Van Strien et al., 2015) and dispersal simulations.

The evaluation and monitoring of *inbreeding and relatedness* can provide valuable information to maximise existing genetic variation and avoid inbreeding depression. This information has been used in captive breeding programs to prevent the detrimental effects of small population size, founder effects, and lack of gene flow (Wright et al., 2021). Various functions can guide the breeding of plants and animals; **gl.grm()** calculates and plots the mean probability of identity of descent across all loci that would result from all the possible crosses of the individuals that were sampled (Fig. 4; Endelman & Jannink, 2012). This information can identify potential pairs of individuals whose crossing might prevent inbreeding.

**Figure 4.**
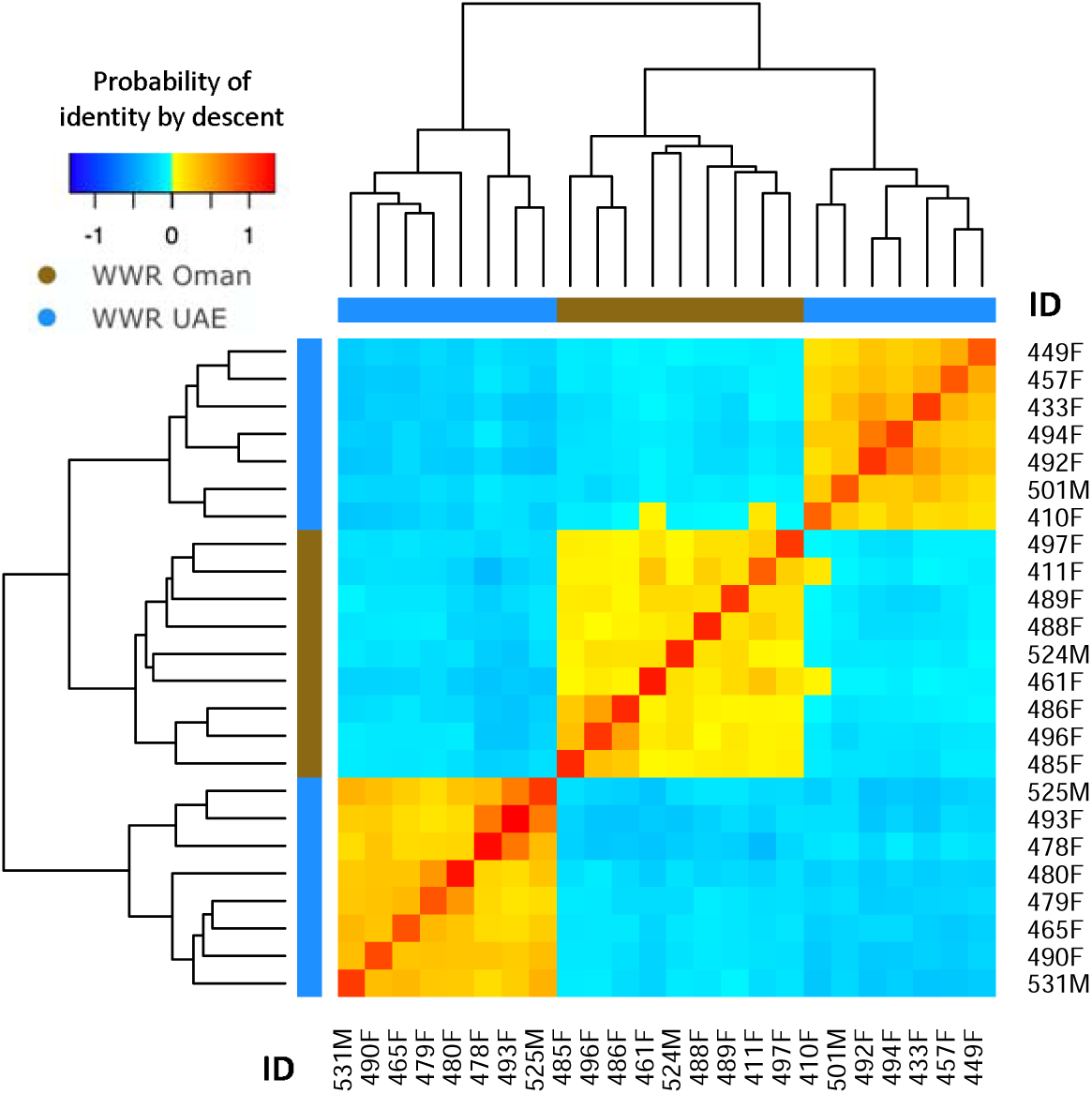
Heatmap of the probabilities of identity by descent (IBD) in which yellow and red colours indicate individuals more related to each other. The identification number of each individual is shown in the margins of the figure, where the last letter denotes whether the individual is male (M) or female (F). This information is being used to guide the captive breeding program of the Arabian oryx at the Al-Wusta Wildlife Reserve in Oman (Al Rawahi *et al*., 2022).

We have developed functions to simplify the process of *running external software* that requires several steps (*a.k.a*. wrapping functions), linking to programs such as Outflank (Whitlock & Lotterhos, 2015), BLAST (Altschul et al., 1990; Altschul et al., 1997), NewHybrids (Anderson & Thompson, 2002), Neestimator2 (Do et al., 2014), STRUCTURE (Pritchard et al., 2000), Clumpp (Jakobsson & Rosenberg, 2007), Distruct (Rosenberg, 2004) and Evanno’s method (Evanno et al., 2005). For example, the latter four programs can be run within dartR using the functions below and results plotted in an interactive map as shown in Fig. 5. Note that while we aimed to facilitate access to resources and analytical tools, the users should remain aware of assumptions and characteristics of such analyses so that they can be run and interpreted properly. We envisage that future version of dartR will continue the development of functions that will facilitate testing of assumption and screening of adequate execution (*e.g*. convergence).

**Figure 5.**
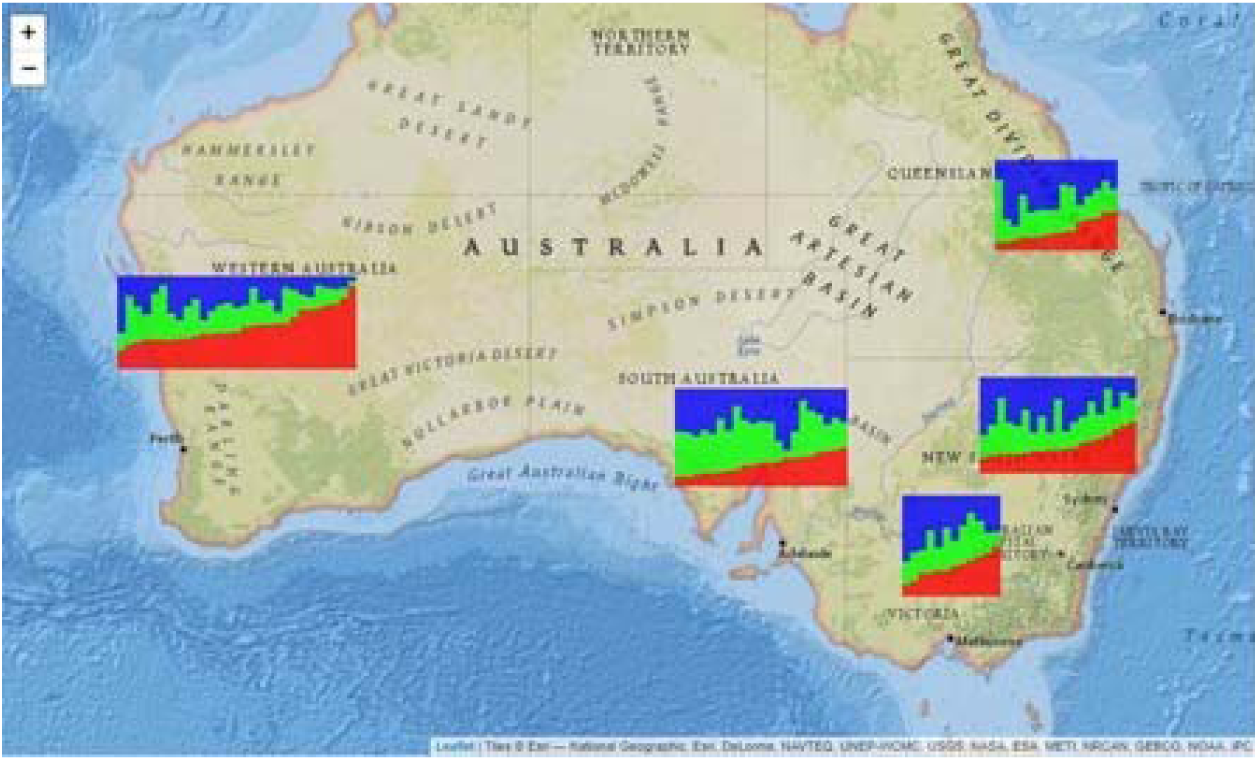
Interactive map showing the results from the software STRUCTURE (Pritchard et al., 2000), using the software Clumpp (Jakobsson & Rosenberg, 2007) to align the results of different independent runs and the software Distruct (Rosenberg, 2004) to display the results graphically. Individuals are shown as vertical bars coloured in proportion to their estimated ancestry within each inferred population (*K*=3). The dataset used in the figure is provided with the package.

**> out_struc <- gl.run.structure(bandicoot.gl, k.range = 2:5, num.k.rep = 10, exec = “∼/structure.exe”, noadmix=FALSE)**

**> out_evanno <- gl.evanno(out_struc)**

**> qmat <- gl.plot.structure(out_struc, k=3, CLUMPP=“∼/CLUMPP.exe”)**

**> gl.map.structure(qmat, bandicoot.gl)**

*Exporting genetic data* to other formats is a common step and one of the most time-consuming and susceptible to errors in the analysis of genetic data. dartR offers 24 functions to export genlight objects to other formats, including FASTA, PLINK and VCF.

*Computer simulations* are powerful tools for understanding complex evolutionary and genetic processes and their relationships to ecological processes and can be used, for example, to predict complex scenarios involving the interaction between evolutionary forces or evaluate the plausibility of alternative hypotheses or, validate and evaluate genetic methods (Hoban et al., 2012). In this second version of dartR, we developed a realistic simulation model that can be parameterised with real-life genetic characteristics such as the number, location, allele frequency and the distribution of fitness effects (selection coefficients and dominance) of loci under selection. In the simulation model recombination is accurately modeled, and it is possible to use real recombination maps as input.

We have also developed a set of internal functions that facilitate the user’s interaction with dartR. For example, the function **gl.install.vanilla.dartR()** installs all required packages for using all the functions available in dartR; and the functions **gl.print.history()** and **gl.play.history()** prints and replays the history of all the analyses performed previously in a genlight object, respectively.

## Concluding remarks

The remarkable recent advances in applied and theoretical genetics offer many novel opportunities to address and better manage rates of biodiversity and ecosystem loss. Notwithstanding this, the list of skills and level of expertise required to integrate novel genomic resources and perform increasingly complex analyses have increased simultaneously. Thus, researchers and stakeholders often struggle to keep up with the various ways to analyse and apply genetic data and to take maximum advantage of them to inform conservation and restoration. We envision that as the number of analyses and their complexity continues to increase, accessible, streamlined and reliable platforms to analyse genetic data, such as dartR, will play a crucial role in translating science into practice.

## Acknowledgements

This project and the scripts that are included were funded in part by a grant from the ACT Priority Investment Fund, the Cooperative Research Network for Murray-Darling Basin Futures and the ARC Linkage Program.

## Author contributions

B.G., A.G. and O.B. conceived the ideas and methodology; B.G., A.G., J.L.M. and C.P. develop the package; J.L.M. led the writing of the manuscript. All authors contributed critically to the drafts and gave final approval for publication.

## Data availability statement

The current version of the dartR package (2.0.3) can be downloaded and installed via CRAN R repository [install.packages(”dartR”)]. The latest development version is hosted on GitHub under: https://github.com/green-striped-gecko/dartR, accompanied by a detailed description of how to install the latest version and a changelog. Errors, feature requests and contributions should be submitted via the GitHub repository.

## References

Al Rawahi, Q., Mijangos, J. L., Khatkar, M. S., Al Abri, M. A., AlJahdhami, M. H., Kaden, J., Gongora, J. (2022). Rescued back from extinction in the wild: past, present and future of the genetics of the Arabian oryx in Oman. Royal Society Open Science, 9(3), 210558.

Altschul, S. F., Gish, W., Miller, W., Myers, E. W., & Lipman, D. J. (1990). Basic local alignment search tool. Journal of molecular biology, 215(3), 403–410.

Altschul, S. F., Madden, T. L., Schäffer, A. A., Zhang, J., Zhang, Z., Miller, W., & Lipman, D. J. (1997). Gapped BLAST and PSI-BLAST: a new generation of protein database search programs. Nucleic Acids Research, 25(17), 3389–3402.

Anderson, E., & Thompson, E. (2002). A model-based method for identifying species hybrids using multilocus genetic data. Genetics, 160(3), 1217–1229.

Andrews, K. R., & Luikart, G. (2014). Recent novel approaches for population genomics data analysis. In: Wiley Online Library.

Bourret, V., Albert, V., April, J., Côté, G., & Morissette, O. (2020). Past, present and future contributions of evolutionary biology to wildlife forensics, management and conservation. Evolutionary Applications, 13(6), 1420–1434.

Breed, M. F., Harrison, P. A., Blyth, C., Byrne, M., Gaget, V., Gellie, N. J., Prowse, T. A. (2019). The potential of genomics for restoring ecosystems and biodiversity. Nature Reviews Genetics, 20(10), 615–628.

Cayuela, H., Rougemont, Q., Prunier, J. G., Moore, J. S., Clobert, J., Besnard, A., & Bernatchez, L. (2018). Demographic and genetic approaches to study dispersal in wild animal populations: A methodological review. Molecular Ecology, 27(20), 3976–4010.

De Meeûs, T. (2018). Revisiting F IS, F ST, Wahlund effects, and null alleles. Journal of Heredity, 109(4), 446–456.

De Meeûs, T., McCoy, K. D., Prugnolle, F., Chevillon, C., Durand, P., Hurtrez-Bousses, S., & Renaud, F. (2007). Population genetics and molecular epidemiology or how to “débusquer la bête”. Infection, Genetics and Evolution, 7(2), 308–332.

Do, C., Waples, R. S., Peel, D., Macbeth, G. M., Tillett, B. J., & Ovenden, J. R. (2014). NeEstimator v2: re-implementation of software for the estimation of contemporary effective population size (Ne) from genetic data. Molecular Ecology Resources, 14(1), 209–214.

Durka, W., Michalski, S. G., Berendzen, K. W., Bossdorf, O., Bucharova, A., Hermann, J. M.,. Kollmann, J. (2017). Genetic differentiation within multiple common grassland plants supports seed transfer zones for ecological restoration. Journal of Applied Ecology, 54(1), 116–126.

Endelman, J. B., & Jannink, J.-L. (2012). Shrinkage estimation of the realized relationship matrix. G3: Genes, Genomes, Genetics, 2(11), 1405–1413.

Evanno, G., Regnaut, S., & Goudet, J. (2005). Detecting the number of clusters of individuals using the software STRUCTURE: a simulation study. Molecular Ecology, 14(8), 2611–2620.

Funk, W. C., McKay, J. K., Hohenlohe, P. A., & Allendorf, F. W. (2012). Harnessing genomics for delineating conservation units. Trends in Ecology & Evolution, 27(9), 489–496.

Garner, B. A., Hand, B. K., Amish, S. J., Bernatchez, L., Foster, J. T., Miller, K. M., Roffler, G. (2016). Genomics in conservation: case studies and bridging the gap between data and application. Trends in Ecology & Evolution, 31(2), 81–83.

Georges, A., Gruber, B., Pauly, G. B., White, D., Adams, M., Young, M. J., Unmack, P. J. (2018). Genomewide SNP markers breathe new life into phylogeography and species delimitation for the problematic short-necked turtles (Chelidae: Emydura) of eastern Australia. Molecular Ecology, 27(24), 5195–5213.

Gruber, B., Unmack, P. J., Berry, O. F., & Georges, A. (2018). dartr: An r package to facilitate analysis of SNP data generated from reduced representation genome sequencing. Molecular Ecology Resources, 18(3), 691–699.

Hoban, S., Bertorelle, G., & Gaggiotti, O. E. (2012). Computer simulations: tools for population and evolutionary genetics. Nature Reviews Genetics, 13(2), 110–122.

Hohenlohe, P. A., Funk, W. C., & Rajora, O. P. (2021). Population genomics for wildlife conservation and management. Molecular Ecology, 30(1), 62–82.

Jakobsson, M., & Rosenberg, N. A. (2007). CLUMPP: a cluster matching and permutation program for dealing with label switching and multimodality in analysis of population structure. Bioinformatics, 23(14), 1801–1806.

Jombart, T., & Ahmed, I. (2011). adegenet 1.3-1: new tools for the analysis of genome-wide SNP data. Bioinformatics, 27(21), 3070–3071.

Kilian, A., Wenzl, P., Huttner, E., Carling, J., Xia, L., Blois, H., Hopper, C. (2012). Diversity arrays technology: a generic genome profiling technology on open platforms. Methods in Molecular Biology, 888(1), 67–89.

McVean, G. (2009). A genealogical interpretation of principal components analysis. PLoS Genet, 5(10), e1000686.

Narum, S. R., Buerkle, C. A., Davey, J. W., Miller, M. R., & Hohenlohe, P. A. (2013). Genotyping-by-sequencing in ecological and conservation genomics. Molecular Ecology, 22(11), 2841–2847.

O’Leary, S. J., Puritz, J. B., Willis, S. C., Hollenbeck, C. M., & Portnoy, D. S. (2018). These aren’t the loci you’e looking for: Principles of effective SNP filtering for molecular ecologists. Molecular Ecology, 27(16), 3193–3206.

Pritchard, J. K., Stephens, M., & Donnelly, P. (2000). Inference of population structure using multilocus genotype data. Genetics, 155(2), 945–959.

R Core Team. (2021). R: A language and environment for statistical computing. R Foundation for Statistical Computing, Vienna, Austria., URL https://www.R-project.org/.

Robertson, B. C., & Gemmell, N. J. (2004). Defining eradication units to control invasive pests. Journal of Applied Ecology, 41(6), 1042–1048.

Rosenberg, N. A. (2004). DISTRUCT: a program for the graphical display of population structure. Molecular Ecology Notes, 4(1), 137–138.

Shafer, A. B., Wolf, J. B., Alves, P. C., Bergström, L., Bruford, M. W., Brännström, I., Ekblom, R. (2015). Genomics and the challenging translation into conservation practice. Trends in Ecology & Evolution, 30(2), 78–87.

Sherwin, W. B., Chao, A., Jost, L., & Smouse, P. E. (2017). Information theory broadens the spectrum of molecular ecology and evolution. Trends in Ecology & Evolution, 32(12), 948–963.

Taylor, H. R., Dussex, N., & van Heezik, Y. (2017). Bridging the conservation genetics gap by identifying barriers to implementation for conservation practitioners. Global Ecology and Conservation, 10, 231–242.

Tigano, A., & Friesen, V. L. (2016). Genomics of local adaptation with gene flow. Molecular Ecology, 25(10), 2144–2164.

Unmack, P. J., Adams, M., Hammer, M. P., Johnson, J. B., Gruber, B., Gilles, A., Georges, A. (2022). Plotting for change: an analytical framework to aid decisions on which lineages are candidate species in phylogenomic species discovery. Biological Journal of the Linnean Society, 135(1), 117–137.

Van Strien, M. J., Holderegger, R., & Van Heck, H. J. (2015). Isolation-by-distance in landscapes: considerations for landscape genetics. Heredity, 114(1), 27–37.

Waples, R. S. (2015). Testing for Hardy–Weinberg proportions: have we lost the plot? Journal of Heredity, 106(1), 1–19.

Weigand, H., & Leese, F. (2018). Detecting signatures of positive selection in non-model species using genomic data. Zoological Journal of the Linnean Society, 184(2), 528–583.

Whitlock, M. C., & Lotterhos, K. E. (2015). Reliable detection of loci responsible for local adaptation: Inference of a null model through trimming the distribution of FST. The American Naturalist, 186(S1), S24–S36.

Wright, B. R., Hogg, C. J., McLennan, E. A., Belov, K., & Grueber, C. E. (2021). Assessing evolutionary processes over time in a conservation breeding program: A combined approach using molecular data, simulations and pedigree analysis. Biodiversity and Conservation, 30(4), 1011–1029.

