## Supplementary Table 1 for "dartR v2: an accessible genetic analysis platform for conservation, ecology, and agriculture"

### 1.1 Supplementary information

**Supplementary Table 1.** Grouping and description of all the functions currently available in dartR.

| Function group | Function name | Description |
| --- | --- | --- |
| Input functions | gl.compliance.check | Checks a genlight object to see if it complies with dartR expectations and amends it to comply if necessary |
|  | gl.read.csv | Reads SNP data from a csv file into a genlight object |
|  | gl.read.dart | Imports DArT data into dartR and converts it into a genlight object |
|  | gl.read.silicodart | Imports presence/absence data from SilicoDArT to genlight format (ploidy=1) |
|  | gl.read.vcf | Converts a vcf file into a genlight object |
| Data manipulation functions | gl.define.pop | Defines a new population in a genlight object for specified individuals |
|  | gl.drop.ind | Removes specified individuals from a genlight object |
|  | gl.drop.loc | Removes specified loci from a genlight object |
|  | gl.drop.pop | Removes specified populations from a genlight object |
|  | gl.edit.recode.ind | Creates or edits individual names, creates a recode_ind file and applies the changes to a genlight object |
|  | gl.edit.recode.pop | Creates or edits a population re-assignment table |
|  | gl.impute | Imputates missing data |
|  | gl.join | Combines two genlight objects |
|  | gl.keep.ind | Removes all but the specified individuals from a genlight object |
|  | gl.keep.loc | Removes all but the specified loci from a genlight object |
|  | gl.keep.pop | Removes all but the specified populations from a genlight object |
|  | gl.make.recode.ind | Creates a proforma recode_ind file for reassigning individual(=specimen) names |
|  | gl.make.recode.pop | Creates a proforma recode_pop_table file for reassigning population names |
|  | gl.merge.pop | Merges two or more populations in a genlight object into one population |
|  | gl.random.snp | Randomly changes the allocation of 0's and 2's in a genlight object |
|  | gl.reassign.pop | Assigns an individual metric as pop in a genlight object |
|  | gl.recalc.metrics | Recalculates locus metrics when individuals or populations are deleted from a genlight object |
|  | gl.recode.ind | Recodes individual labels in a genlight object |
|  | gl.recode.pop | Recodes population assignments in a genlight object |
|  | gl.rename.pop | Renames a population in a genlight object |
|  | gl.subsample.loci | Subsamples n loci from a genlight object and return it as a genlight object |
| Reporting functions | gl.diagnostics.hwe | Provides descriptive stats and plots to diagnose potential problems with Hardy-Weinberg proportions |
|  | gl.diagnostics.sim | Compares simulations against theoretical expectations |
|  | gl.report.bases | Reports summary of base pair frequencies |
|  | gl.report.callrate | Reports summary of Call Rate for loci or individuals |
|  | gl.report.diversity | Calculates diversity indexes for SNPs |
|  | gl.report.hamming | Calculates the pairwise Hamming distance between DArT trimmed DNAsequences |
|  | gl.report.heterozygosity | Reports observed, expected and unbiased heterozygosities and FIS (inbreeding coefficient) by population or by individual from SNP data |
|  | gl.report.hwe | Reports departure from Hardy-Weinberg proportions |
|  | gl.report.ld.map | Calculates pairwise linkage disequilibrium in SNPs mapped to a reference genome |
|  | gl.report.ld | Calculates pairwise population based Linkage Disequilibrium across all loci using the specified number of cores |
|  | gl.report.locmetric | Reports summary of the slot $other$loc.metrics |
|  | gl.report.maf | Reports minor allele frequency (MAF) for each locus in a SNP dataset |
|  | gl.report.monomorphs | Reports monomorphic loci |
|  | gl.report.overshoot | Reports loci for which the SNP has been trimmed from the sequence tag along with the adaptor |
|  | gl.report.pa | Reports private alleles (and fixed alleles) per pair of populations |
|  | gl.report.parent.offspring | Identifies putative parent offspring within a population |
|  | gl.report.rdepth | Reports summary of Read Depth for each locus |
|  | gl.reporte.reproducibility | Reports summary of RepAvg (repeatability averaged over both alleles foreach locus) or reproducibility (repeatability of the scores for fragment presence/absence) |
|  | gl.report.secondaries | Reports loci containing secondary SNPs in sequence tags and calculatesnumber of invariant sites |
|  | gl.report.sexlinked | Identifies loci that are sex linked |
|  | gl.report.taglength | Reports summary of sequence tag length across loci |
| Filtering functions | gl.filter.allna | Filters loci that are all NA across individuals and/or populations with all NA across loci |
|  | gl.filter.callrate | Filters loci or specimens in a genlight object based on callrate |
|  | gl.filter.hamming | Filters loci based on pairwise Hamming distance between sequence tags |
|  | gl.filter.heterozygosity | Filters individuals with average heterozygosity greater than a specified upper threshold or less than a specified lower threshold |
|  | gl.filter.hwe | Filters loci that show significant departure from Hardy-Weinberg Equilibrium |
|  | gl.filter.locmetric | Filters loci on the basis of numeric information stored in other$loc.metrics in a genlight object |
|  | gl.filter.maf | Filters loci on the basis of minor allele frequency (MAF) in a genlight object |
|  | gl.filter.monomorphs | Filters monomorphic loci, including those with all NAs |
|  | gl.filter.overshoot | Filters loci for which the SNP has been trimmed from the sequence tag along with the adaptor |
|  | gl.filter.pa | Filters loci that contain private (and fixed alleles) between twopopulations |
|  | gl.filter.parent.offspring | Filters putative parent offspring within a population |
|  | gl.filter.rdepth | Filters loci based on counts of sequence tags scored at a locus (read depth) |
|  | gl.filter.reproducibility | Filters loci in a genlight object based on average repeatability of alleles at a locus |
|  | gl.filter.secondaries | Filters loci that represent secondary SNPs in a genlight object |
|  | gl.filter.sexlinked | Filters loci that are sex linked |
|  | gl.filter.taglength | Filters loci in a genlight object based on sequence tag length |
| Exploration and visualisation functions | gl.grm | Calculates an identity by descent matrix |
|  | gl.grm.network | Represents a genomic relationship matrix (GRM) as a network |
|  | gl.map.interactive | Creates an interactive map (based on latlon) from a genlight object |
|  | gl.map.structure | Maps a STRUCTURE plot using a genlight object |
|  | gl.pcoa | Ordination applied to genotypes in a genlight object (PCA), in an fdobject, or to a distance matrix (PCoA) |
|  | gl.pcoa.plot | Bivariate or trivariate plot of the results of an ordination generated using gl.pcoa() |
|  | gl.plot.heatmap | Represents a distance matrix as a heatmap |
|  | gl.plot.network | Represents a distance or dissimilarity matrix as a network |
|  | gl.plot.structure | Plots a STRUCTURE analysis using a genlight object |
|  | gl.smearplot | Smear plot of SNP or presence/absence (SilicoDArT) data |
| Genetic variation functions | gl.alf | Calculates allele frequency of the first and second allele for each loci |
|  | gl.amova | Performs AMOVA using genlight data |
|  | gl.basic.stats | Calculates basic statistics for each loci (Hs, Ho, Fis etc.) |
|  | gl.fst.pop | Calculates a pairwise Fst values for populations in a genlight object |
|  | gl.He | Estimates expected Heterozygosity |
|  | gl.Ho | Estimates observed Heterozygosity |
|  | gl.hwe.pop | Performs Hardy-Weinberg tests over loci and populations |
|  | gl.percent.freq | Generates percentage allele frequencies by locus and population |
|  | gl.report.diversity | Calculates diversity indexes for SNPs |
|  | gl.report.heterozygosity | Reports observed, expected and unbiased heterozygosities and FIS (inbreeding coefficient) by population or by individual from SNP data |
|  | gl.report.pa | Reports private alleles (and fixed alleles) per pair of populations |
|  | gl.test.heterozygosity | Tests the difference in heterozygosity between populations taken pairwise |
|  | is.fixed | Tests if two populations are fixed at a given locus |
| Population identification functions | gl.assign.mahalanobis | Assign an individual of unknown provenance to population based on Mahalanobis distance |
|  | gl.assign.pa | Eliminates populations as possible source populations for an individual of unknown provenance, using private alleles |
|  | gl.assign.pca | Assign an individual of unknown provenance to population based on PCA |
|  | gl.collapse | Collapses a distance matrix by amalgamating populations with pairwise fixed difference count less that a threshold |
|  | gl.fdsim | Estimates the rate of false positives in a fixed difference analysis |
|  | gl.fixed.diff | Generates a matrix of fixed differences and associated statistics for populations taken pairwise |
|  | gl.nhybrids | Creates an input file for the program NewHybrids and runs it if NewHybrids is installed |
|  | gl.pcoa | Ordination applied to genotypes in a genlight object (PCA), in an fdobject, or to a distance matrix (PCoA) |
|  | gl.pcoa.plot | Bivariate or trivariate plot of the results of an ordination generated using gl.pcoa() |
|  | gl.run.structure | Runs a STRUCTURE analysis using a genlight object |
|  | gl.tree.nj | Outputs an nj tree to summarize genetic similarity among populations |
| Dispersal and gene flow functions | gl.assign.pa | Eliminates populations as possible source populations for an individual of unknown provenance, using private alleles |
|  | gl.costdistances | Calculates cost distances for a given landscape (resistance matrix) |
|  | gl.dist.ind | Calculates a distance matrix for individuals defined in a genlight object |
|  | gl.dist.pop | Calculates a distance matrix for populations with SNP genotypes in a genlight object |
|  | gl.fst.pop | Calculates a pairwise Fst values for populations in a genlight object |
|  | gl.genleastcost | Performs least-cost path analysis based on a friction matrix |
|  | gl.ibd | Performs isolation by distance analysis |
|  | gl.propShared | Calculates a similarity (distance) matrix for individuals on the proportion of shared alleles |
| Inbreeding and relatedness functions | gl.filter.parent.offspring | Filters putative parent offspring within a population |
|  | gl.grm | Calculates an identity by descent matrix |
|  | gl.grm.network | Represents a genomic relationship matrix (GRM) as a network |
|  | gl.report.parent.offspring | Identifies putative parent offspring within a population |
| Functions to use external software | gl.blast | Aligns nucleotides sequences against those present in a target database using blastn |
|  | gl.evanno | Creates an Evanno plot from an STRUCTURE run object |
|  | gl.LDNe | Estimates effective population size using the Linkage Disequilibrium method based on NeEstimator (V2) |
|  | gl.nhybrids | Creates an input file for the program NewHybrids and runs it if NewHybrids is installed |
|  | gl.outflank | Identifies loci under selection per population using the outflank method of Whitlock and Lotterhos (2015) |
|  | gl.plot.structure | Plots a STRUCTURE analysis using a genlight object |
|  | gl.run.structure | Runs a STRUCTURE analysis using a genlight object |
| Exporting functions | gi2gl | Converts a genind object into a genlight object |
|  | gl.write.csv | Writes out data from a genlight object to csv file |
|  | gl2bayescan | Converts a genlight object into a format suitable for input to Bayescan |
|  | gl2demerelate | Creates a dataframe suitable for input to package Demerelate from a genlight object |
|  | gl2eigenstrat | Converts a genlight object into eigenstrat format |
|  | gl2fasta | Concatenates DArT trimmed sequences and outputs a FASTA file |
|  | gl2faststructure | Converts a genlight object into faststructure format (to run faststructure elsewhere) |
|  | gl2gds | Converts a genlight object into gds format |
|  | gl2genalex | Converts a genlight object into a format suitable for input to genalex |
|  | gl2genepop | Converts a genlight object into genepop format |
|  | gl2geno | Converts a genlight object to geno format from package LEA |
|  | gl2gi | Converts a genlight object to genind object |
|  | gl2hiphop | Converts a genlight objects into hiphop format |
|  | gl2phylip | Creates a Phylip input distance matrix from a genlight (SNP) object |
|  | gl2plink | Converts a genlight object into PLINK format |
|  | gl2related | Converts a genlight object to format suitable to be run with Coancestry |
|  | gl2sa | Converts a genlight object to the format used in the SNPassoc package |
|  | gl2sfs | Converts a genlight object into a sfs input file |
|  | gl2shp | Converts a genlight object to ESRI shapefiles or kml files |
|  | gl2snapp | Converts a genlight object to nexus format suitable for phylogenetic analysis by SNAPP (via BEAUti) |
|  | gl2structure | Converts a genlight object to STRUCTURE formated files |
|  | gl2svdquartets | Converts a genlight object to nexus format PAUP SVDquartets |
|  | gl2treemix | Converts a genlight object to a treemix input file |
|  | gl2vcf | Converts a genlight object into vcf format |
| Simulation functions | gl.sim.create.dispersal | Creates a dispersal file as input for the function gl.sim.WF.run |
|  | gl.sim.emigration | Simulates emigration between populations |
|  | gl.sim.ind | Simulates individuals based on the allele frequencies provided via a genlight object. |
|  | gl.sim.mutate | Simulates mutations within a genlight object |
|  | gl.sim.offspring | Simulates a specified number of offspring based on alleles provided by potential father(s) and mother(s) |
|  | gl.sim.WF.run | Runs Wright-Fisher simulations |
|  | gl.sim.WF.table | Creates the reference table for running gl.sim.WF.run |
| Internal functions | gl.check.verbosity | Checks the current global verbosity |
|  | gl.install.vanilla.dartR | Installs all required packages for using all functions available in dartR |
|  | gl.list.reports | Prints dartR reports saved in tempdir |
|  | gl.load | Loads an object from compressed binary format produced by gl.save() |
|  | gl.play.history | Replays the history and applies it to a genlight object |
|  | gl.print.history | Prints history of a genlight object |
|  | gl.print.reports | Prints dartR reports saved in tempdir |
|  | gl.save | Saves an object in compressed binary format for later rapid retrieval |
|  | gl.select.colors | Selects colors from one of several palettes and output as a vector |
|  | gl.select.shapes | Selects shapes from the base R shape palette and outputs as a vector |
|  | gl.set.verbosity | Sets the default verbosity level |
